# BOLD Contrast Response Characteristics of Aberrant Voxels with Bilateral Visual Population Receptive Fields in Human Albinism

**DOI:** 10.1101/2024.05.26.595603

**Authors:** Ethan J. Duwell, Erica N. Woertz, Jedidiah Mathis, Joseph Carroll, Edgar A. DeYoe

**Author notes:** Corresponding Author: Edgar DeYoe, Department of Radiology, Medical College of Wisconsin, 8701 Watertown Plank Road, Milwaukee, WI 53226.

## Abstract

Albinism is an inherited disorder characterized by disrupted melanin production in the eye, and often in the skin and hair. This retinal hypopigmentation is accompanied by pathological decussation of many temporal retinal afferents at the optic chiasm during development, ultimately resulting in partially superimposed representations of opposite visual hemifields in each cortical hemisphere. Within these aberrant regions of hemifield overlap, individual voxels have been shown to have bilateral, dual population receptive fields (pRFs) responding to roughly mirror-image locations across the vertical meridian. Nonetheless, how these two conflicting inputs combine to determine a voxel’s response to image contrast is still unknown. To address this, we stimulated the right and left hemifields with separately controlled sinusoidal gratings, each having a variety of contrasts (0, 8, 20, 45, 100%), and extracted voxel-wise BOLD response amplitudes to each contrast combination in visual areas V1-V3. We then compared voxels’ responses to each hemifield stimulated individually with conditions when both hemifields were stimulated simultaneously. We hypothesized that simultaneous stimulation of the two pRF components will result in either a suppressive or facilitative interaction. However, we found that BOLD responses to simultaneous stimulation appeared to reflect simple summation of the neural activity from the individual hemifield conditions. This suggests that the superimposed opposite hemifield representations do not interact. Thus, dual pRFs in albinism likely reflect two co-localized, but functionally independent populations of neurons each of which respond to a single hemifield. This finding is commensurate with psychophysical studies which have shown no clear perceptual interaction between opposite visual hemifields in human albinism.

## INTRODUCTION

The human visual system is typically organized contralaterally such that the right visual hemifield is represented in the left cortical hemisphere and vice versa. This organization is established by the normal patterns of retinal afferent decussation at the optic chiasm whereby the temporal afferents continue ipsilaterally and the nasal afferents cross to the contralateral hemisphere. Afferents from equivalent retinal locations in each eye therefore join at the chiasm, synapse in the LGN contralateral to their visual field input, and ultimately reach the same cortical regions in V1 such that fMRI imaging voxels at each cortical location respond to the same contralateral field location in each eye. However, while imaging voxels respond macroscopically to both eyes, individual layer-four cells in V1 receive monocular input and are segregated into alternating ocular dominance columns.

In albinism, the typical contralateral organization is corrupted due to a significant proportion of the temporal retinal afferents from each eye which decussate pathologically at the optic chiasm. Consequently, in subjects with albinism, each cortical hemisphere receives input from both the contra- and ipsilateral visual fields, ultimately resulting in partially superimposed representations of opposite hemifields in each hemisphere (Hoffmann, Tolhurst, Moore, & Morland, 2003; Morland, Baseler, Hoffmann, Sharpe, & Wandell, 2001). The extent of these aberrant hemifield overlap zones varies across individual subjects with albinism depending on the severity of miswiring (Hoffmann et al., 2003). Individual fMRI voxels within the aberrant zones were demonstrated to have bilateral, dual pRFs positioned at roughly mirror image locations across the vertical meridian (Ahmadi, Herbik, Wagner, Kanowski, Thieme, & Hoffmann, 2019; Carvalho, Invernizzi, Ahmadi, Hoffmann, Renken, & Cornelissen, 2020; Duwell, Woertz, Mathis, Carroll, & DeYoe, 2021). However, the cellular and columnar organization of these aberrant cortical zones is still not known in human albinism, and it is unclear whether the opposing hemifield inputs interact in any way. Consequently, voxels with dual pRFs in albinism could potentially reflect either a single population of neurons which receive dual hemifield input, two co-localized populations of neurons responding to each respective hemifield that interact in some way, or two co-localized but functionally separate populations of neurons that do not interact. The sole electrophysiological study on non-human albino primates suggests that ocular dominance columns in these zones are replaced by hemifield columns (Guillery, Hickey, Kaas, Felleman, Debruyn, & Sparks, 1984), and that these hemifield columns do not interact. This hemifield columnar pattern was also observed in a recent ultra-high resolution 7T fMRI study of a human subject with achiasma (Olman, Bao, Engel, Grant, Purington, Qiu, Schallmo, & Tjan, 2018). However, columnar organization has yet to be established in human albinism.

Psychophysical studies in human albinism have shown no perceptual interaction between opposite hemifields suggesting that the cortically superimposed hemifield representations may remain functionally separate at the cellular level (Hoffmann, Seufert, & Schmidtborn, 2007; Klemen, Hoffmann, & Chambers). Similarly, a recent study of achiasma showed that psychophysical contrast detection thresholds in each hemifield were not affected by contrast presented to the opposite hemifield, and that stimulating opposite hemifields simultaneously did not result in BOLD fMRI responses indicative of a neural interaction between the opposite hemifield inputs (Bao, Purington, & Tjan, 2015). Instead, BOLD responses to both hemifields stimulated simultaneously were consistent with a simple summation of neural activity from the two individual hemifields. However, similar fMRI contrast response experiments have not been performed in human albinism and it’s possible that dual pRF voxels in albinism may respond differently to simultaneous stimulation of opposite hemifields. To address this, we stimulated the right and left hemifields both individually and simultaneously with separately controlled sinusoidal gratings, each having a variety of contrasts (0, 8, 20, 45, 100%) and extracted voxels’ BOLD response amplitudes to each contrast combination in visual areas V1-V3 using linear regression. These experiments and analyses were inspired by Bao et al. and the original fMRI contrast experiments by Boynton et al. (Bao et al., 2015; Boynton, Demb, Glover, & Heeger, 1999). We assessed voxels’ BOLD contrast responses to each hemifield individually and compared these to responses when contrast was presented to both hemifields simultaneously. Our results suggest that the underlying neural responses to each hemifield individually add linearly during simultaneous stimulation, and that the opposing hemifield inputs therefore do not interact. Consequently, as in achiasma, dual pRF voxels in albinism likely reflect separate, co-localized populations of non-interacting cells which each respond to input from a single visual hemifield.

## METHODS

### Subjects

Four subjects with albinism (3 females, 1 male; aged 23-36 years) with minimal nystagmus and three control subjects with no prior ocular or cortical pathology (2 females, 1 male; aged 28-34 years) were recruited for this experiment. All subjects in our albinism cohort participated in our previous pRF and cortical magnification studies (Duwell et al., 2021; Woertz, Wilk, Duwell, Mathis, Carroll, & DeYoe, 2020). Genetic information and demographics for subjects with albinism were described in our previous studies but are repeated for this subset of subjects in **Table 1** below. Fixation stability in these subjects was also previously characterized (Duwell et al., 2021; Woertz et al., 2020). This study was in accordance with the Declaration of Helsinki and approved by the Institutional Review Board of the Medical College of Wisconsin. All subjects were provided written informed consent after explanation of the nature and possible risks and benefits of the study.

**Table 1:** Genetics and Demographics for Subjects with Albinism.

| Subject | ID* | Age (y) | Sex | BCVA | Albinism Subtype^ | Mutations |
| --- | --- | --- | --- | --- | --- | --- |
| A1 | JC_10227 | 23 | F | 20/40 (OU) | OCA2 | OCA2 c.1327G>A; p.V443I<br>TYR c.575C>A; p.S192Y |
| A2** | JC_0492 | 36 | F | 20/25 (OU) | OCA1 | TYR c.1147G>A; p.D383N<br>TYR c.1217C>T; p.P406L |
| A3** | JC_0493 | 28 | F | 20/20- (OU) | OCA1 | TYR c.1147G>A; p.D383N<br>TYR c.1217C>T; p.P406L |
| A4 | JC_10093 | 19 | M | 20/100 +2 (OD)<br>20/80 +2 (OS) | OA | GPR143 c.346T>G; p.C116G<br>TYR c.1205G>A; p.R402Q (hom) |

\*ID used in Wilk *et al.* (2014) & Wilk *et al.* (2017)
\*\*Subjects are sisters
^OCA=oculocutaneous albinism; OA=ocular albinism (X-linked)
(hom) = homozygous

### fMRI Visual Stimuli

Note: Detailed descriptions of the retinotopic ring and wedge stimuli, visual area mapping techniques, and image processing methods for the monocular, hemifield eccentricity data and binocular, full-field pRF modeling data used in this study are described in our recent study of aberrant visual pRFs in albinism (Duwell et al., 2021). This study utilizes those same data for the albinism cohort. Monocular, hemifield eccentricity data, and full field pRF modeling data were acquired for control subjects using identical methods.

Contrast stimuli for fMRI were presented on a back-projection screen mounted on the MRI head coil. A BrainLogics BLMRDP-A05 MR digital projector was used with a ViSaGe MKII visual stimulus generator (Cambridge Research Instruments Ltd, Rochester, UK) in conjunction with custom MATLAB software. All stimuli were photopic, presented binocularly on a uniform gray background, and subtended a maximum of 20° eccentricity. A schematic depicting several frames of our hemifield-contrast stimulation task is depicted in **Figure 1**. Contrast stimuli were composed of counter-phase flickering sinusoidal gratings angled at −45 and 45 degrees in the left and right hemifields respectively. Orthogonal angles were presented in opposite hemifields in hopes of increasing the possibility of a hemifield interaction in the albinism group. As illustrated in **Figure 1**, contrast presentation trials followed a boxcar design with 8 s. ON periods followed by 8 s. OFF periods of grey background. During each trial, separate gratings of either 0, 8, 20, 45, or 100 percent contrast appeared in each hemifield. Each of the 25 possible hemifield-contrast pairs were presented once per run for a total of 416 s. per run (including before and after periods). Runs were split into 5, 80 s. blocks, each containing 5 trials. During each run, contrast in one hemifield stepped sequentially from 0-100 for trials within each block while the opposite hemifield contrast remained constant for trials within each block but stepped sequentially from 0-100 across blocks. The hemifield displaying within- vs. across-block contrast increments alternated right vs. left on each run. To control fixation and attention, subjects were instructed to fixate on a black, circular, central fixation point (radius = 0.3 degrees) and to press a button whenever it blinked white. Fixation point blinks were 200 ms. and occurred pseudo-randomly every 1-4 s.

**Figure 1:**
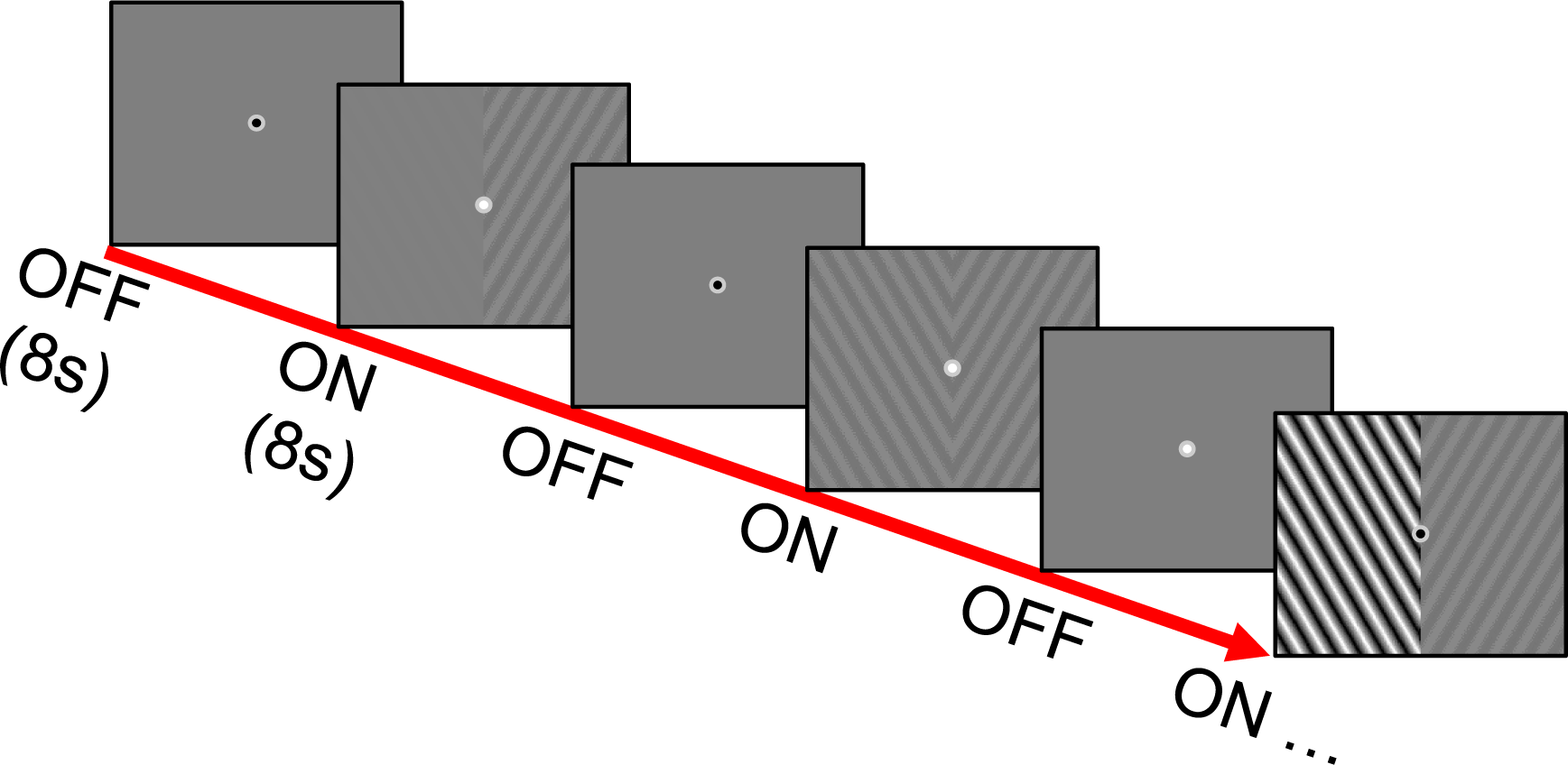
Schematic of the hemifield-contrast stimulation paradigm. Example frames depict 8 s. ON and OFF periods. During ON periods, counterphase flickering sinusoidal gratings of either 0, 8, 20, 45, or 100% contrast appeared in the right and left hemifields. Contrast was controlled independently for each hemifield and each of the 25 possible hemifield-contrast combinations was presented once per run. Grey background was displayed during the 8 s. OFF periods. Subjects were instructed to fixate on the circular fixation marker (center of each frame) and press a button whenever it blinked white.

8 runs were collected for each subject. All subjects with albinism completed scanning in a single session and runs presenting the two sequential orders were interleaved. Due to scan time and scheduling constraints, two of the control subjects completed their runs across two separate sessions. Runs presenting the two sequential orders were still interleaved and split evenly across two scanning sessions.

### fMRI Acquisition

Scan acquisition parameters for the BOLD contrast response experiments in this study were intentionally identical to those used in our recent pRF modeling study in albinism (Duwell et al., 2021). MRI scans were obtained at the Medical College of Wisconsin using a 3.0 Tesla General Electric Signa Discovery 750 MRI system equipped with a custom 32-channel RF/gradient head coil. BOLD fMRI images were acquired with a T2*-weighted gradient-echo EPI pulse sequence (TE = 25 ms, TR = 2 s, FA = 77°). The 96 x 96 acquisition matrix (Fourier interpolated to 128 x 128) had frequency encoding in the right-left axial plane, phase encoding in the anterior-posterior direction and slice selection in the axial direction. The FOV was 240 mm and included 29 axial slices covering the occipital lobe and adjacent portions of the temporal and parietal lobes with a slice thickness of 2.5 mm, yielding a raw voxel size of 1.875 x 1.875 x 2.5 mm. For anatomical scans, a T1-weighted, spoiled gradient recalled at steady state (SPGR), echo-planar pulse sequence was used (TE = 3.2 ms, TR = 8.2 ms, FA = 12°) with a 256 x 224 acquisition matrix (Fourier interpolated to 256 x 256). The FOV was 240 mm, and 180 slices with a slice thickness of 1.0 mm, yielding voxel sizes of 0.938 × 0.938 × 1.0 mm^3^. The SPGR scans were subsequently resampled to 1.0 mm^3^. A sync pulse from the scanner at the beginning of each run triggered the onset of visual stimuli.

### Analysis Software

All fMRI data were analyzed using the AFNI/SUMA software package (version: AFNI_19.1.11, https://afni.nimh.nih.gov/) (Cox, 1996). Surface models were produced from the high resolution SPGR images using the ‘recon-all’ function in Freesurfer (version 5.1.0 and 5.3.0, http://surfer.nmr.mgh.harvard.edu/). All, pRF and contrast response function modeling was performed using custom software in MATLAB (2017b and 2019a).

### fMRI Pre-processing

fMRI pre-processing was performed using a script generated by AFNI’s afni_proc.py command and occurred in the following order: reconstruction, removal of before and after periods, slice timing correction, alignment and volume registration, smoothing, masking, scaling, regression. Before and after periods were removed using AFNI 3dTcat, and slice time shift correction was then performed using AFNI 3dTshift. Functional EPI runs were aligned to the anatomical SPGRs collected during the previous monocular hemifield expanding ring and pRF modeling experiments (Duwell et al., 2021) to allow voxel-wise comparison between the contrast response and pRF modeling data. SUMA cortical surface models from those previous experiments were also used. Rigid body alignment and volume registration were performed using AFNIs align_epi_anat.py and 3dVolreg respectively. During volume registration, volumes in each run were registered to the volume in that series determined to be the ‘minimum outlier’ by AFNI 3dToutcount. This volume was also used as the base EPI for aligning each functional run to the anatomical scan. The alignment and volume registration transformations were computed separately, concatenated, and then applied together. The 6 registration parameters from 3dVolreg were used to compute motion magnitude time series which served as motion regressors later in the pipeline. Each run was then smoothed with a 3.75 mm (full width at half max) Gaussian kernel using AFNI 3dMerge, and then brain-masked using 3dAutomask. After masking, the timeseries data were then scaled to range from 0-200 with a mean of 100 and were subsequently demeaned such that final EPI timeseries express percentage of the mean. Finally, the scaled runs were concatenated, and the regression analysis was performed using AFNI 3dDeconvolve. The regression model included a third order polynomial detrend and the six motion parameter timeseries (concatenated across runs) as baseline model regressors. Volumes preceding motion magnitudes >0.3 mm and volumes in which >5% of voxels within the brain were computed as outliers (determined by AFNI 3doutcount) were censored in the regression analysis. 25 regressors of interest were also included corresponding to the 25 different hemifield-contrast combinations presented throughout the experiment. Each regressor consisted of a separate stimulus timing file containing the stimulus onset times of the respective hemifield-contrast conditions in the concatenated dataset. We used AFNI’s BLOCK option as our regression basis function with a block length of 8 s. to account for the trial presentation length and a regressor unit height of 1. Our regressor and timeseries scaling parameters were chosen such that the beta coefficient outputs for each regressor express voxels’ response amplitudes to each condition in units of % BOLD change. Finally, we used AFNI 3dresample to match the voxel grid coordinates in the statistical output map to the corresponding pRF modeling and retinotopic datasets to which they were aligned.

### Voxel Selection

To select voxels with the highest quality responses, subjects’ fMRI contrast response datasets were percentile thresholded based on the full f-stat output of the linear regression model. Voxels with f-stats in the 90^th^ percentile within subjects’ brain-masked EPI datasets were included. This percentile method allowed for variability in data quality across subjects while maintaining a proportional level of threshold rigor. On average, this amounted to corrected p-value thresholds of 2.5E^−06^ and 2.4E^−04^ in the albinism and control groups respectively. Individual subjects’ equivalent Bonferroni-corrected p-value thresholds are listed on the right-hand column in **Figure 2**. Results presented in this study were also further restricted to voxels falling within our V1-V3 visual area ROIs. Furthermore, to control for eye and hemifield input, we selected subgroups of voxels which responded exclusively to either the contralateral hemifield in both eyes or to opposite hemifields in the contralateral eye. The latter only occur in the albinism group due to retinocortical miswiring. Voxels’ eye and hemifield inputs were determined independently in previous monocular, individual hemifield, expanding ring experiments thresholded at a correlation coefficient of 0.45. For more information on these methods see (Duwell et al., 2021).

**Figure 2:**
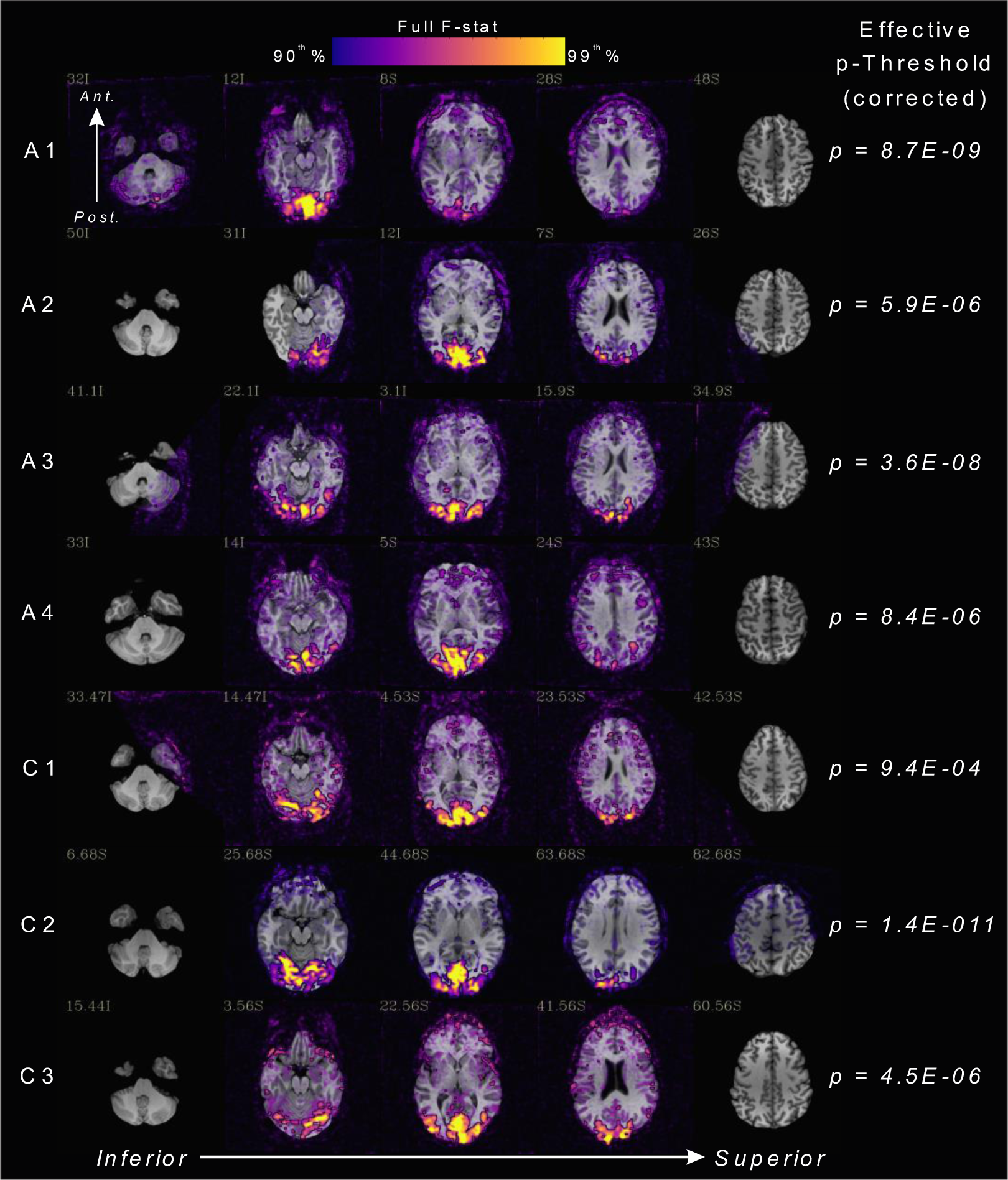
fMRI activation maps for the hemifield-contrast stimulation experiment in subjects with albinism (rows A1-A4) and controls (rows C1-C3). Colored overlays (ranging from purple to yellow) represent the full F-stat percentile from the regression analysis ranging from the 90th-99th percentile in each subject (see color key). Subjects’ respective Bonferroni corrected p-value thresholds equivalent to 90th percentile are listed on the right. Task data are superimposed on axial slices of subject’s anatomical SPGR images.

### Assessing Contrast Response Summation

Beta weights expressing voxels’ BOLD contrast response amplitudes in each of the 25 hemifield/contrast conditions were extracted and categorized with respect to contrasts presented to the contra- and ipsilateral hemifields relative to each voxel’s cortical hemisphere. To assess the extent to which voxels’ responses to simultaneous stimulation of opposite hemifields reflect the sum of responses to the contra- and ipsilateral hemfields alone, for each voxel, we computed the sum of responses to all pairs of non-zero contrasts presented to the contralateral and ipsilateral hemifields individually:

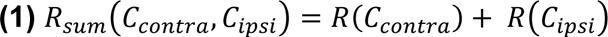

*C_contra_* and *C_ipsi_* are the contrasts presented to the contra- and ipsilateral hemifields, *R(C_contra_)* and *R(C_ipsi_)* are the empirical BOLD responses in the individual hemifield stimulation conditions, and *R_sum_* is simply the sum of the two responses.

However, the relationship between the BOLD response and the relative underlying level of neural activity is known to be non-linear. Best estimates suggest that the BOLD response is roughly proportional to the sum of the underlying neural activity raised to the 0.5 power (Bao et al., 2015). To account for this, we also computed a predicted BOLD response to simultaneous stimulation of opposite hemifields for voxels in the hemifield overlap zones in albinism which incorporates this non-linear relationship between neural activity and the BOLD response. This predicted response assumed that the neural activity in response to the contra- and ipsilateral hemifield conditions adds linearly during simultaneous stimulation and that the voxel’s BOLD response is proportional to the sum of neural activity raised to the 0.5 power. As responses to each hemifield/contrast combination are measured within the same voxel, hemodynamics are constant and we therefore assumed that differences in BOLD responses across conditions reflect differences in the underlying neural activity. Given our assumed relationship between neural activity and the BOLD response, it follows that relative changes in BOLD response between conditions must reflect relative changes in neural activity raised to the 0.5 power, and conversely, that relative changes in neural activity reflect changes in BOLD response squared:

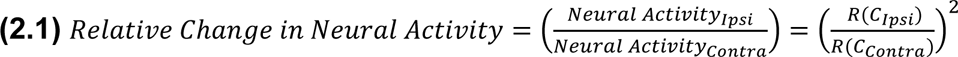

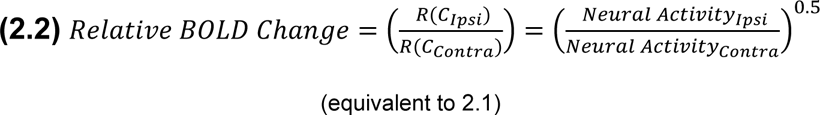

Given our assumption that neural activities in the respective contra- and ipsilateral hemifield stimulation conditions simply add during simultaneous stimulation of opposite hemifields, the relative change in neural activity between the contralateral hemifield condition alone and simultaneous stimulation of opposite hemifields can be expressed as:

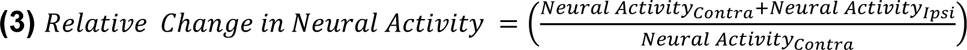

Given **(2.2)** the relative change in BOLD response between the two conditions must then be:

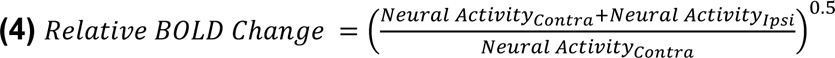

As neural activity is assumed to be proportional to the BOLD responses squared, equation **(4)** can also be expressed in terms of the BOLD responses to the contra and ipsilateral stimulation conditions (*R*(*C*_*Contra*_) and *R*(*C*_Ipsi_)) as follows:

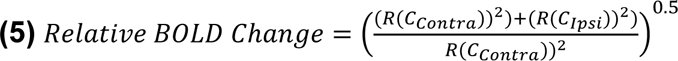

Consequently, given a voxel’s measured BOLD responses to the contra- and ipsilateral conditions individually, the response to the contra- and ipsilateral contrasts presented simultaneously can be predicted by multiplying the empirical BOLD response to the contralateral condition alone (*R(C_contra_)*) by the relative BOLD change in (**5**) above:

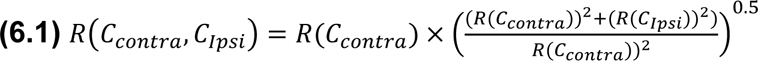

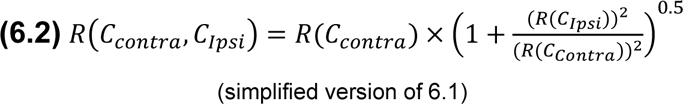

*C_contra_* and *C_ipsi_* are, again, the contrasts presented to the contra- and ipsilateral hemifields. *R(C_contra_) and R(C_ipsi_)* are the empirical BOLD responses in the individual hemifield stimulation conditions, and *R(C_contra_,C_ipsi_)* is the predicted BOLD response to simultaneous *C_contra_* and *C_ipsi_* stimulation assuming that neural activities from the individual hemfifield conditions sum, and the BOLD response is proportional to the summed neural activity raised to the 0.5 power.

## RESULTS

### Contrast Responses for Normal and Aberrant Voxels in Albinism and Controls

**Figure 2** displays fMRI task activation data from the hemifield contrast experiment overlaid on the SPGR anatomical images for each subject with albinism (rows A1-A4) and controls (rows C1-C3). Colored overlays represent the whole-brain, full *f*-stat percentile from the regression analysis in each subject ranging from the 90th-99th percentile. Equivalent p-value thresholds corresponding to each subject’s 90^th^ % full *f-*stat are displayed on the right. All subjects showed highly significant BOLD responses to our hemifield-contrast stimulation task in occipital visual cortex. Coverage of V1-V3 visual area ROIs was comparable and nearly complete in all subjects with albinism and controls.

BOLD contrast response data are presented in **Figures 3A (control)** and **3B (albinism)** for voxels that respond conventionally to a single pRF in the contralateral hemifield. **Figures 3C** and **3D** present contrast response data for dual-pRF voxels in albinism which respond aberrantly to inputs from opposite hemifields in the contralateral eye. Different colors represent mean BOLD response to each of 5 contrasts (0, 8, 20, 45, or 100%) presented in the opposite hemifield (color code in **Figure 3**, upper right). Normal voxels in both the control (**3A**) and albinism groups (**3B**) showed little systematic shift in their response functions as contrast was increased in the ipsilateral hemifield. For aberrant voxels in albinism though, contrast response curves in the contralateral hemifield clearly shifted systematically with increasing contrast in the ipsilateral hemifield (**3C**) and vice versa (**3D**). However, BOLD response amplitudes to stimulation of the contralateral hemifield alone (green data points in **3D**) appeared more robust (higher amplitude) on average than corresponding aberrant responses to the ipsilateral hemifield alone (green data points in **3C**). This effect is also apparent in the greater baseline shifts induced by contralateral stimulation in **3D** vs. ipsilateral stimulation in **3C.** It is noteworthy that for the aberrant dual pRF voxels in **3C** and **3D**, there also appeared to be a compression of the asymptotic maximum contrast response and a reduction in slope across visual areas V1-V3. This reduction in slope suggests a loss in contrast response gain across visual areas. This effect is not apparent for normally wired voxels in either controls (**3A**) or subjects with albinism in (**3B**).

**Figure 3:**
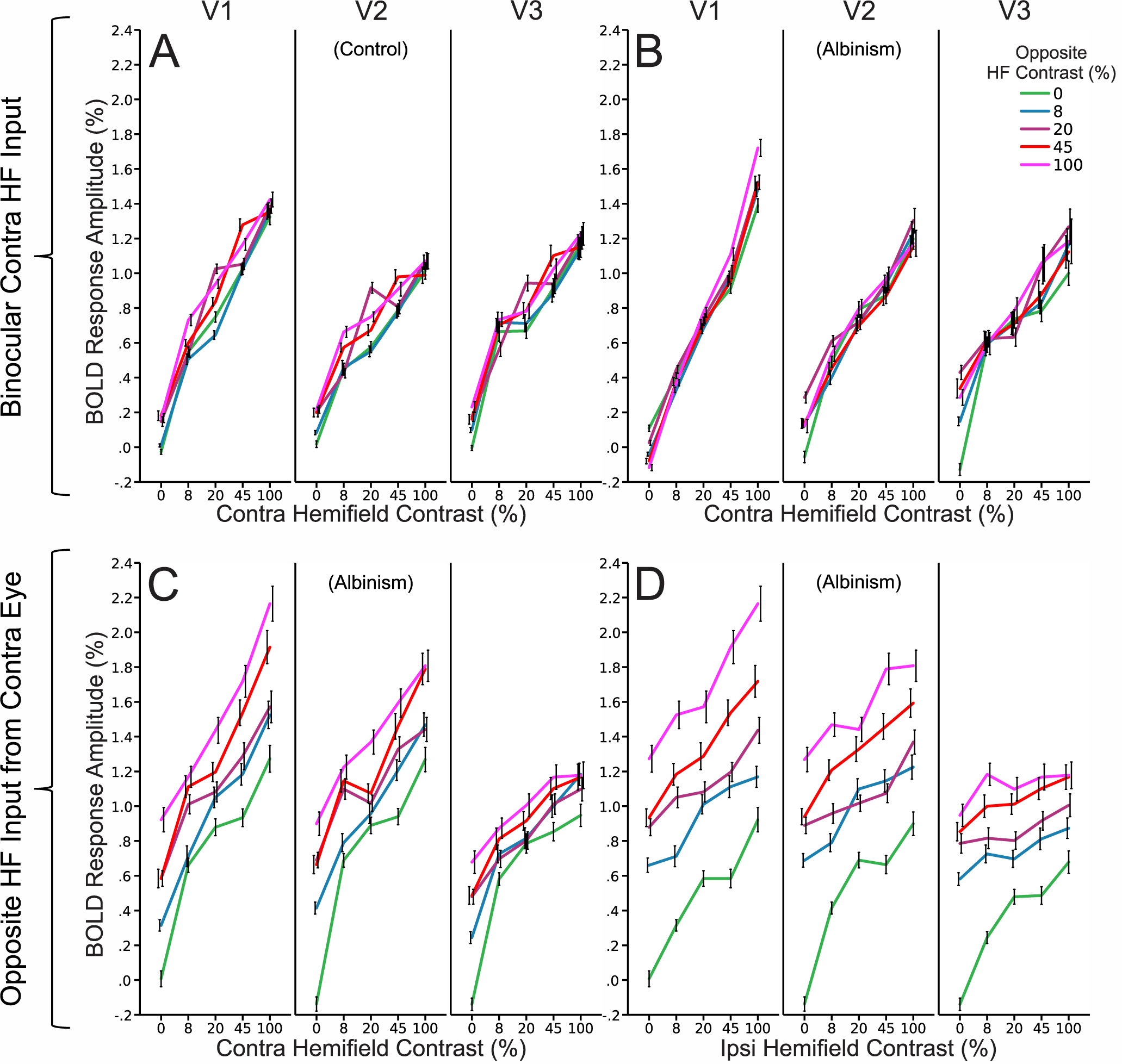
Group BOLD contrast response data for normal voxels which respond binocularly to the contralateral hemifield in controls (A) and albinism (B) and for aberrant voxels in albinism which respond to opposite hemifields in the contralateral eye (C and D) displayed separately for V1, V2, and V3 (columns). Colored line plots represent BOLD responses with 0, 8, 20, 45, and 100% contrast in the opposite hemifield (color code in upper right). Data in (A) and (B) plot contralateral hemifield responses (x axes) as modulated by the ipsilateral hemifield (colors). (C) and (D) plot contra- and ipislateral hemifield responses respectively (x axes) as modulated by the ipsi- and contralateral hemifields (colors). (C and D are the same data plotted two different ways). Data represent the mean response amplitudes for all voxels pooled across subjects in each group. Error bars represent the 95% confidence interval. Contra = Contralateral, Ipsi = Ipsilateral.

Contrast response data in **Figure 3C** and **3D** are displayed for each individual subject with albinism in **Figures 4A** and **4B** respectively. There was some variability in overall response amplitude across subjects, and ipsilateral contrast responses were especially robust in subjects A1 and A4. However, the observations in group data above in **3C** and **3D** are clearly visible in each individual subject’s data in **4A** and **4B**.

**Figure 4:**
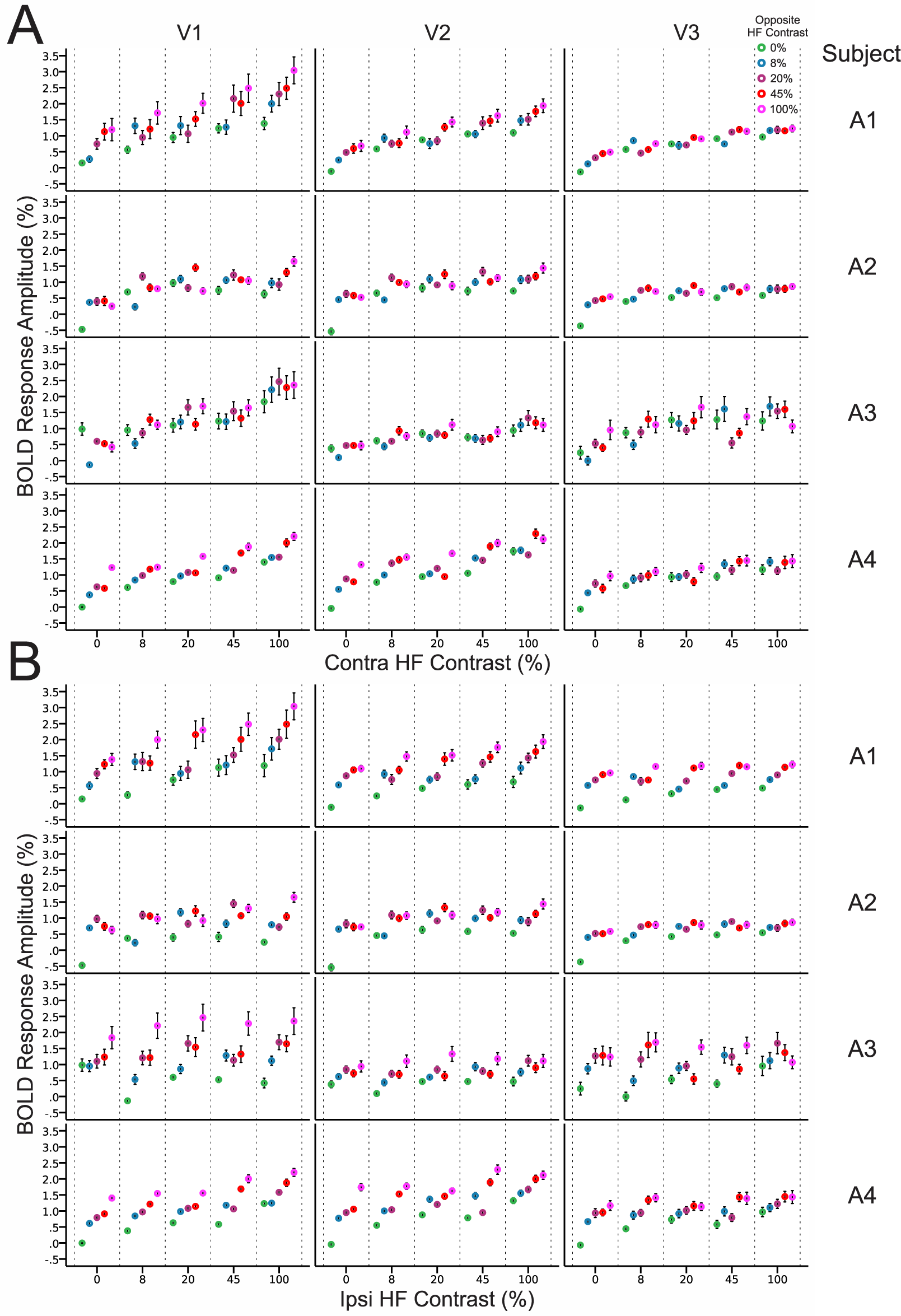
BOLD contrast responses to the contralateral (A) and ipsilateral hemifields (B) for aberrant voxels driven by opposite hemifields in the contralateral eye for individual subjects with albinism (rows A1-A4). (A and B are the same data plotted two different ways.) Colors represent responses with 0, 8, 20, 45 and 100% contrast in the opposite hemifield (color code in upper right). Data are displayed separately for V1, V2 and V3 (columns). Values and error bars represent the mean and 95% confidence intervals across all aberrant voxels in each subject with albinism. Contra = Contralateral, Ipsi = Ipsilateral.

### Assessing BOLD Contrast Response Summation for Aberrant Voxels in Albinism

Focusing on the dual pRF voxels in albinism, **Figure 5** illustrates the extent to which BOLD responses to simultaneous stimulation of opposite hemifields reflect the sum of responses to each hemifield individually. Green curves represent empirical responses to simultaneous stimulation of opposite hemifields; whereas, blue curves were predicted by linear summation (**Equation 1**) of the BOLD responses to each hemifield stimulated alone. The red curves are responses predicted by summation of *neural activity* from the two individual hemifield stimulation conditions assuming that the BOLD response is proportional to the square root of the summed neural activity (**Equation 6.2**). Each plot contains empirical and predicted responses to each contrast in the contralateral hemifield (x axis) and each row of plots display responses with 0, 8, 20, 45 and 100% contrast in the ipsilateral hemifield respectively. In most cases, the empirical responses to simultaneous hemifield stimulation (green) were remarkably close to those predicted by summation of neural activity from the individual hemifield conditions combined with the known nonlinearity between neural and BOLD responses (red). These two mean curves coincided almost exactly in most conditions. They were especially close in lower contrast stimulation conditions and in V2-V3. As reported by Bao et al. in achiasma (Bao et al., 2015), the computed sum of the BOLD responses to each hemifield individually (blue) was consistently much higher than the response to both hemifields stimulated simultaneously. However, for some high contrast conditions (45-100%), simultaneous stimulation produced responses that were best predicted as the linear sum of the separate hemifield responses (blue), especially in V1.

**Figure 5:**
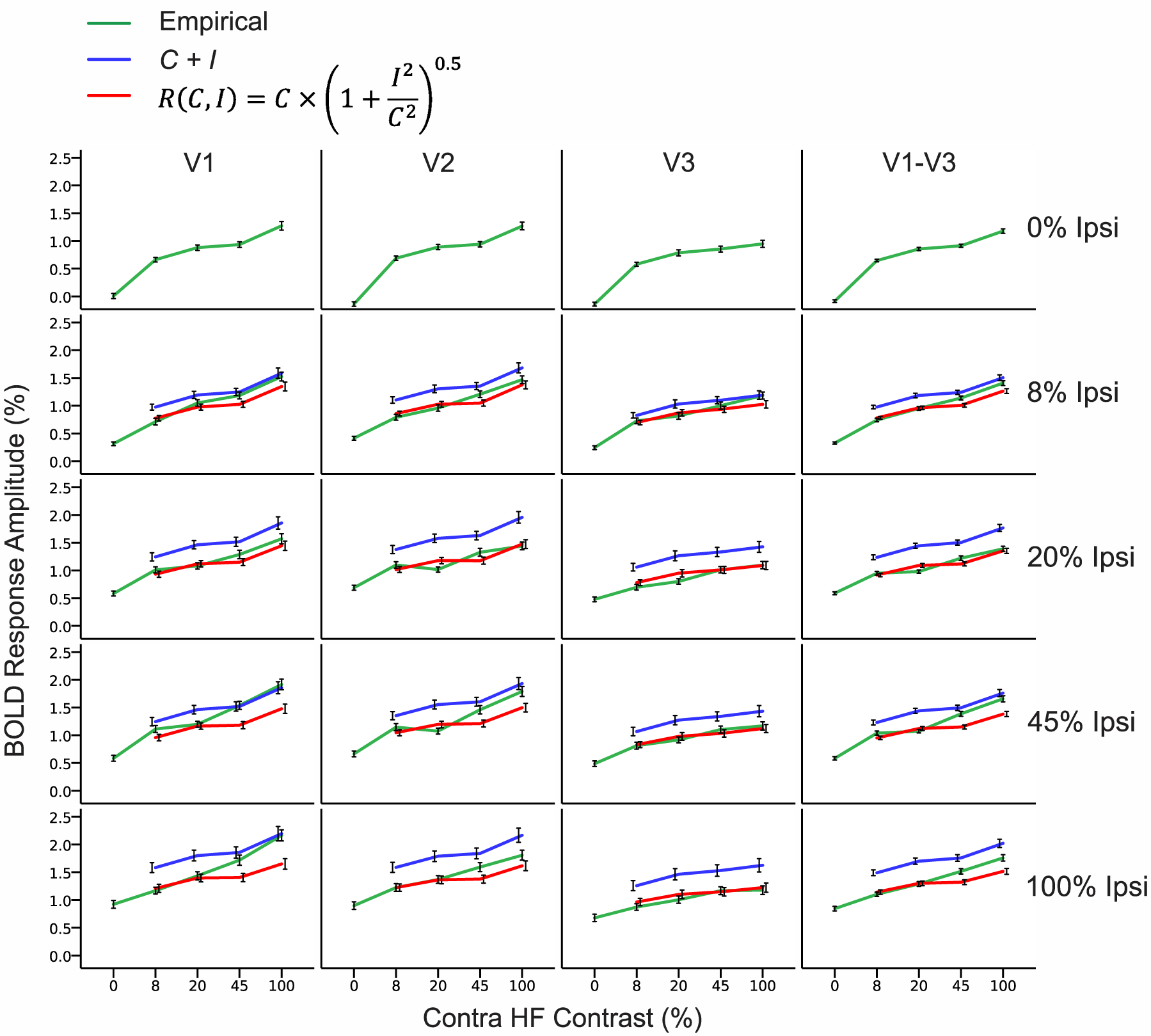
Empirical BOLD contrast responses to simultaneous stimulation of opposite hemifields (green) vs. the sum of individual hemifield condition responses (blue) and responses predicted by the summed ‘neural responses’ from the individual hemifield conditions assuming the BOLD response is proportional to the neural response raised to the 0.5 power (red). Data are plotted for V1-V3 separately and combined (columns) for aberrant voxels in albinism which respond to opposite hemifields in the contralateral eye. Contralateral hemifield contrast is plotted along the x axes and data in each row plot conditions with 0, 8, 20, 45, and 100% contrast in the ipsilateral hemifield. Values represent the mean BOLD amplitudes for all voxels pooled across subjects and error bars are the 95% confidence interval. Empirical responses (green) closely follow those predicted by neural summation of the individual hemfield responses combined with the known non-linear BOLD/neural activity relationship (red), but sometimes fall between this prediction and pure BOLD summation (blue).

Overall, responses to simultaneous hemifield stimulation were clearly not greater than the linear sum of the individual hemifield responses and were also not less than responses predicted by simple neural summation combined with the known nonlinearity between neural activity and the BOLD response. This suggests that there were neither facilitative nor suppressive interactions between the conflicting opposite hemifield inputs in albinism. Instead, these data are consistent with two co-localized but functionally separate populations of neurons within each voxel which respond to the contra- and ipsilateral fields respectively, and whose activities consequently sum during simultaneous stimulation of both hemifields.

## DISCUSSION

In this study we examined the BOLD contrast response characteristics of aberrant voxels in human albinism which respond to both the contra- and ipsilateral visual hemifields in the contralateral eye. Previous studies have shown that retinocortical miswiring in albinism results in partial cortical superposition of opposite hemifield representations in visual cortex such that single imaging voxels in these aberrant zones have two visual pRFs positioned roughly at mirror image locations across the vertical meridian (Ahmadi et al., 2019; Carvalho et al., 2020; Duwell et al., 2021; Hoffmann et al., 2003). However, while the opposite hemifield representation topographies are clearly superimposed at the macroscopic level and within individual voxels in albinism, it was still unclear whether the two conflicting hemifield inputs interact in any way. Consequently, voxels with dual visual pRFs in albinism could potentially reflect either a single population of neurons which receive dual hemifield input, two co-localized populations of neurons responding to each respective hemifield that interact in some way, or two co-localized but functionally separate populations of neurons that do not interact. In the former two possibilities, simultaneous stimulation of opposite hemifields would likely result in a response interaction between the two incongruous hemifield inputs; whereas, in the latter scenario, stimulating both hemifields should result in a simple summation of neural activity. To test for potential interactions between the opposite hemifield representations in albinism, we measured the BOLD contrast responses of aberrant voxels in V1-V3 to a variety of grating contrasts in the contra- and ipsilateral hemifields individually and in combination. We hypothesized that simultaneous stimulation of the opposite hemifield inputs would result in BOLD response amplitudes consistent with either a facilitative or suppressive interaction. As expected, voxels in albinism which responded to opposite hemifields in the contralateral eye during our previous retinotopic mapping experiments also responded systematically to contrasts presented in both hemifields. Conversely, normal voxels in albinism and controls which responded binocularly to the contralateral hemifield only responded robustly to contralateral contrasts. Consistent with previous reports in achiasma (Bao et al., 2015), dual pRF responses to simultaneous stimulation of opposite hemifields were significantly less than the sum of their responses to each hemifield alone (**Figure 5**). However, responses to simultaneous hemifield stimulation were typically very close to those predicted by summation of the *neural activity* from the individual hemifield conditions assuming that the BOLD response is roughly proportional to the square root of the underlying neuronal response (as measured by previous studies (Bao et al., 2015)). Together, these results suggest that there are not facilitative or suppressive neuronal interactions between the conflicting hemifield inputs in albinism, and that voxels with dual pRFs likely reflect two co-localized but functionally separate populations of cells which each respond to a single visual location. This is consistent with the only electrophysiological study of columnar organization in albino non-human primates (Guillery et al., 1984), and psychophysical studies which have found no significant perceptual interactions between opposite hemifields in human albinism (Hoffmann et al., 2007; Klemen et al.).

### BOLD Contrast Responses in Albinism and Controls

As illustrated in **Figure 3**, normal voxels in both albinism and control groups responded only to the contralateral hemifield of each eye; whereas, aberrant dual pRF voxels responded to both hemifields of the contralateral eye. Accordingly, dual pRF voxels responded systematically to contrast in both the contra- and ipsilateral hemifields. Moreover, this was observed in all subjects of the albinism group. This systematic effect suggests that dual pRF responses to the ipsilateral hemifield are a biological consequence of retinocortical miswiring. However, the inputs to dual pRF voxels from the two opposing hemifields were not entirely equivalent. Responses to a contrast level presented to the contralateral hemifield tended to be stronger than responses to the same stimulus in the ipsilateral hemifield. This could potentially reflect a systematic imbalance in the number of cells responding to each hemifield or an asymmetry in their relative response strengths. However, further investigation in a larger albinism sample is necessary to establish whether this is a reproducible effect.

### Visual Area Differences in BOLD Contrast Response

BOLD responses to the ipsilateral hemifield were observed in V1, V2, and V3 (**Figures 3-4**) indicating, as in previous studies, that the aberrant ipsilateral representation is propagated up the visual hierarchy (Duwell et al., 2021; Kaule, Wolynski, Gottlob, Stadler, Speck, Kanowski, Meltendorf, Behrens-Baumann, & Hoffmann, 2014; Wolynski, Kanowski, Meltendorf, Behrens-Baumann, & Hoffmann, 2010). Interestingly, the range of contrast response amplitudes to both the contra- and ipsilateral hemifields appeared to vary systematically with visual area with the asymptotic maximum contrast response decreasing systemically with hierarchical level (**Figures 3C-D**, and **4A-B**). This was also accompanied by a reduction in slope potentially indicating differences in contrast response gain across visual areas. This phenomenon was not observed for normal voxels receiving binocular contralateral field input (**Figure 3A-B)**. The biological relevance of this observation is still unclear at this point and this effect needs to be confirmed in a larger sample. However, these contrast response differences across visual areas could potentially reflect changes in cortico-cortical connections between the various visual areas within the hemifield overlap zones or altered integration of inputs moving up the hierarchy.

### Summation of Hemifield Responses During Simultaneous Stimulation

As observed by Bao et al. in achiasma (Bao et al., 2015), the BOLD response amplitudes of aberrant voxels in albinism to both hemifields simultaneously were consistently less than the sum of voxels’ responses to the same contrasts presented to each hemifield individually (**Figure 5**). However, response amplitudes to simultaneous hemifield stimulation closely followed those predicted by summation of neural activity from the individual hemifield conditions assuming the BOLD signal is proportional to the summed neural activity raised to the 0.5 power. This power-law relationship between neural activity and BOLD response amplitude is, on average, what has been reported previously; although, the precise exponent value varies across studies (Bao et al., 2015). In most cases, responses of the aberrant voxels to both hemifields simultaneously closely followed those predicted by summation of the neural activities from the individual hemifield conditions in conjunction with this nonlinearity assumption. This was especially true at lower contrast values and within visual areas V2 and V3 (**Figure 5**). For whatever reason, at higher contrasts (45-100%) in V1, responses to simultaneous stimulation were closer to the sum of the individual hemifield responses. However, as the precise power-law relationship is known to vary slightly across studies, it is likely that optimizing the exponent used in our equation (**6.2**) for this particular dataset would result in a closer prediction for these higher contrast responses. Nonetheless, responses to simultaneous hemifield stimulation were clearly not greater than the sum of the individual hemifield responses or less than those predicted by simple summation of neural activity in light of the typical non-linearity between neural activity and the BOLD response. Overall, these results strongly suggest there are neither facilitative nor suppressive interactions between the partially superimposed opposite hemifield representations in V1-V3 in albinism.

### Limitations and Caveats

One clear limitation of this study is that we only tested for response interactions between opposite hemifields in albinism with respect to a single stimulus property: grating contrast. Our results indicate that the opposite hemifield inputs in albinism do not interact with respect to contrast response amplitude in V1-V3; however, this does not necessarily preclude the possibility of interactions with respect to other stimulus features such as spatial frequency, angle, color, etc. Future studies should investigate potential BOLD and psychophysical interactions between opposite hemifields in albinism with respect to other visual stimulus parameters.

In addition, this study was conducted on a relatively small sample of four subjects with albinism. The contrast responses and BOLD summation patterns we observed in this sample were consistent in the aberrant voxels across these subjects. However, as albinism is a heterogenous condition, these effects could conceivably vary across phenotypic subtypes. In the future, these experiments should be replicated in a larger more heterogeneous albinism sample.

### Implications for Cortical Organization in Albinism

The lack of any clear facilitative or suppressive interactions between the conflicting hemifield inputs in albinism implies that voxels with bilateral dual pRFs reflect two co-localized populations of functionally independent cells which each respond to a single visual field location. This result is consistent with the “True Albino” hemifield column organization and single cell response characteristics observed by Guillery et al. in the only non-human albino primate study, and with psychophysical studies in human albinism which have shown no significant perceptual interactions between opposite hemifields (Guillery et al., 1984; Hoffmann et al., 2007; Klemen et al.). Our results are also consistent with those observed in similar experiments by Bao et al. (2015) in achiasma, suggesting further similarity in aberrant cortical organization between these two populations.

However, the lack of interaction between the hemifield inputs in V1, V2, and V3, also implies that there must be altered cortico-cortical connections, lateral inhibition, and convergence properties across the visual hierarchy in human albinism. We speculate that this may, in some way, underlie the apparent differences in contrast gain and response range across visual areas which we observed in the aberrant, but not the normally wired voxels in **Figure 3**. In normal individuals, monocular cells within ocular dominance columns in V1 eventually converge onto binocular cells in subsequent layers and in higher visual areas such as V2 and V3. These binocular cells and the ubiquitous suppressive lateral connections between ocular dominance columns give rise to known physiological and perceptual interactions between the respective ocular inputs such as response normalization and binocular rivalry (Lee, Blake, & Heeger, 2007; Moradi & Heeger, 2009). If ocular dominance columns are indeed replaced by hemifield columns within the hemifield overlap zones in human albinism and there is also no interaction between the respective hemifield inputs in either V1, V2 or V3, then normal lateral inhibition and convergence across the visual hierarchy must have systematically changed within these zones. If true, this almost certainly has implications for visual function and perception. Consequently, although previous psychophysical studies have not shown direct interactions between the cortically superimposed hemifield locations, future studies focusing on potential consequences of altered lateral inhibition or altered convergence of bottom-up inputs in higher visual areas may reveal perceptual consequences of retino-cortical miswiring in human albinism.

## CONCLUSIONS

In this study, we measured the BOLD contrast response properties of aberrant dual pRF voxels in subjects with albinism versus typical single pRF voxels in albinism and controls across visual areas V1-V3. Aberrant voxels in albinism respond to roughly mirror-symmetric locations in both the contra and ipsilateral hemifields. To test for potential interactions between the conflicting hemifield inputs, we measured voxels’ BOLD responses to a variety of grating contrasts presented to each hemifield individually and in combination. Similar to reports for achiasma (Bao et al., 2015), responses of aberrant dual pRF voxels to simultaneous hemifield stimulation were consistently less than the sum of their responses to each hemifield alone. However, this likely reflects nonlinearities in the BOLD response which typically is proportional to the square root of the underlying neural activity. When this non-linear relationship between neural activity and the BOLD response was taken into account, voxels’ responses to simultaneous hemifield stimulation were accurately predicted by the sum of the neural activity evoked by separate hemifield stimulation. This suggests that superimposed opposite hemifield representations in albinism do not interact, and that voxels with dual pRFs in albinism therefore reflect two co-localized populations of functionally independent cells which each respond to a single hemifield location.

## ACKNOWLEDGEMENTS

We want to acknowledge Steve Jankowski for his tremendous work as our MR technologist, and Erin Curran for her invaluable role in subject recruitment. We also want to acknowledge Andrew Nencka for providing invaluable support while deriving predictive equations relating BOLD signal amplitudes to underlying neural activity. Research reported in this publication was supported by the National Eye Institute, the National Institute of General Medical Sciences, and the National Center for Advancing Translational Sciences of the National Institutes of Health under award numbers TL1TR001437, T32GM080202, T32EY014537, P30EY001931, and R01EY024969. This investigation was conducted in a facility constructed with support from Research Facilities Improvement Program, Grant Number C06RR016511, from the National Center for Research Resources, National Institutes of Health. The content is solely the responsibility of the authors and does not necessarily represent the official views of the National Institutes of Health. This work was also supported by Vision for Tomorrow.

## Contributions

Edgar DeYoe and Ethan Duwell designed the following experiments and visual stimuli. MR image acquisition was performed by Jedidiah Mathis and Ethan Duwell. Ethan Duwell also designed and implemented the image pre-processing pipeline and post-processing analyses. The figures and manuscript were developed by Ethan Duwell with the assistance of Edgar DeYoe, Erica Woertz, Joseph Carroll.

